# Neural crest cell biology shapes lizard skull evolution across evolutionary time scales

**DOI:** 10.1101/2025.07.22.665935

**Authors:** Quentin Horta-Lacueva, Tobias Uller, Morris Flecks, Mariam Gabelaia, Christy Anna Hipsley, Martin Kirchner, Johannes Müller, Nathalie Feiner

**Author notes:** Department of Biology, Lund University, Ekologihuset, Kontaktvägen 10, 223 62 Lund, Sweden.

## Abstract

The vertebrate skull originates from two embryonic lineages, the mesoderm and the neural crest, offering a unique framework to study how developmental mechanisms connect phenotypic variation and evolutionary diversification. Using 3D geometric morphometrics, we analysed skull shape variation in lacertid lizards. Mesoderm- and neural crest-derived bones formed distinct, conserved modules at both micro- and macroevolutionary scales. In the common wall lizard (*Podarcis muralis*), rapid evolution of skull shape under sexual selection was primarily driven by neural crest-derived bones. While the primary axis of shape divergence in *P. murali*s aligned with a major axis of variation across lacertids, neural crest-derived bones exhibited slower evolutionary rates and lower morphological disparity than mesodermal-derived bones. We propose that this discrepancy between the role of the neural crest for skull evolution on micro- and macroevolution reflects constraints imposed by neural crest cell biology: although developmental plasticity enables rapid, correlated responses under sexual selection, pleiotropy may limit long-term evolvability of neural crest-derived skull regions.

**Teaser text:** The bones of the vertebrate skull come from two developmental sources: the mesoderm and the neural crest. This dual origin allows to study how development influences evolution. Using 3D models of lizard skulls—including a species with exaggerated male traits linked to the neural crest—we examined patterns of skull variation. We found that neural crest-derived bones contribute to rapid changes driven by sexual selection. However, across different species, these same skull regions evolve more slowly and show less variation. This suggests that while neural crest cells may constrain long-term evolution because of their wide influence, they can also enable fast adaptations. The developmental biology of a trait therefore shapes its evolution.

## Introduction

Developmental processes structure the morphological variation available to selection (Alberch, 1989; Uller *et al*., 2024). If developmental processes themselves remain conserved, evolutionary adaptation and diversification will be facilitated in some directions (i.e., ‘lines of least resistance’), reflecting phenotypic variants that are both readily generated and adaptive (Schluter, 1996; Tejero-Cicuéndez *et al*., 2023; Rohner & Berger, 2025). Persistent developmental bias may explain why the major axis of phenotypic variation among individuals or between populations of a species is commonly aligned with divergence between species (Jablonski, 2020; Tejero-Cicuéndez *et al*., 2023; Stansfield & Parsons, 2024; Tsuboi *et al*., 2024; Rohner & Berger, 2025). Furthermore, developmental processes contribute to macroevolutionary patterns of diversification. For example, models of tooth development improve the explanatory power of comparative studies in primates and rodents (Kavanagh *et al*., 2007; Machado *et al*., 2023). However, in the absence of a developmental explanation for why phenotypic variation should be structured in a particular way, the processes underlying major evolutionary trends remain challenging to establish.

Structures with more than one developmental origin are valuable to test the impact of developmental bias on evolution. The vertebrate skull is one such structure because it develops from two different germ layers: the mesoderm and the neural crest, the latter derived from the ectoderm (Figure 1; 11, 12). Bones that are mesoderm- or neural crest-derived are partitioned within the skull in a configuration that is largely conserved across vertebrates (Fabbri *et al*., 2017; Teng *et al*., 2019). This dual developmental origin of the skull may make bones of the same origin vary consistently together, imposing a certain structure on morphological variation (Wilson *et al*., 2021). In contrast to the mesoderm, neural crest cells are transient, vertebrate-specific cells that migrate to different parts of the embryo, including the head, and differentiate into a wide range of cell types (e.g., bone, pigmentary, endocrine, mesenchymal, neuronal and glial cells). That neural crest cells migrate and differentiate into many other cell types in addition to those that make up the skull has been proposed to generate covariance between morphological, behavioural and physiological traits (Wilkins *et al*., 2014; Sánchez-Villagra *et al*., 2016; Parsons *et al*., 2020), thereby exercising additional influence over variation and evolution in skull shape. Overall, such differences between mesoderm- and neural crest-derived skull bones should generate structured patterns of covariance (i.e., modularity) that influences morphological adaptation and diversification on both micro- and macroevolutionary time scales (Wilson *et al*., 2021; Goswami *et al*., 2023).

**Figure 1.**
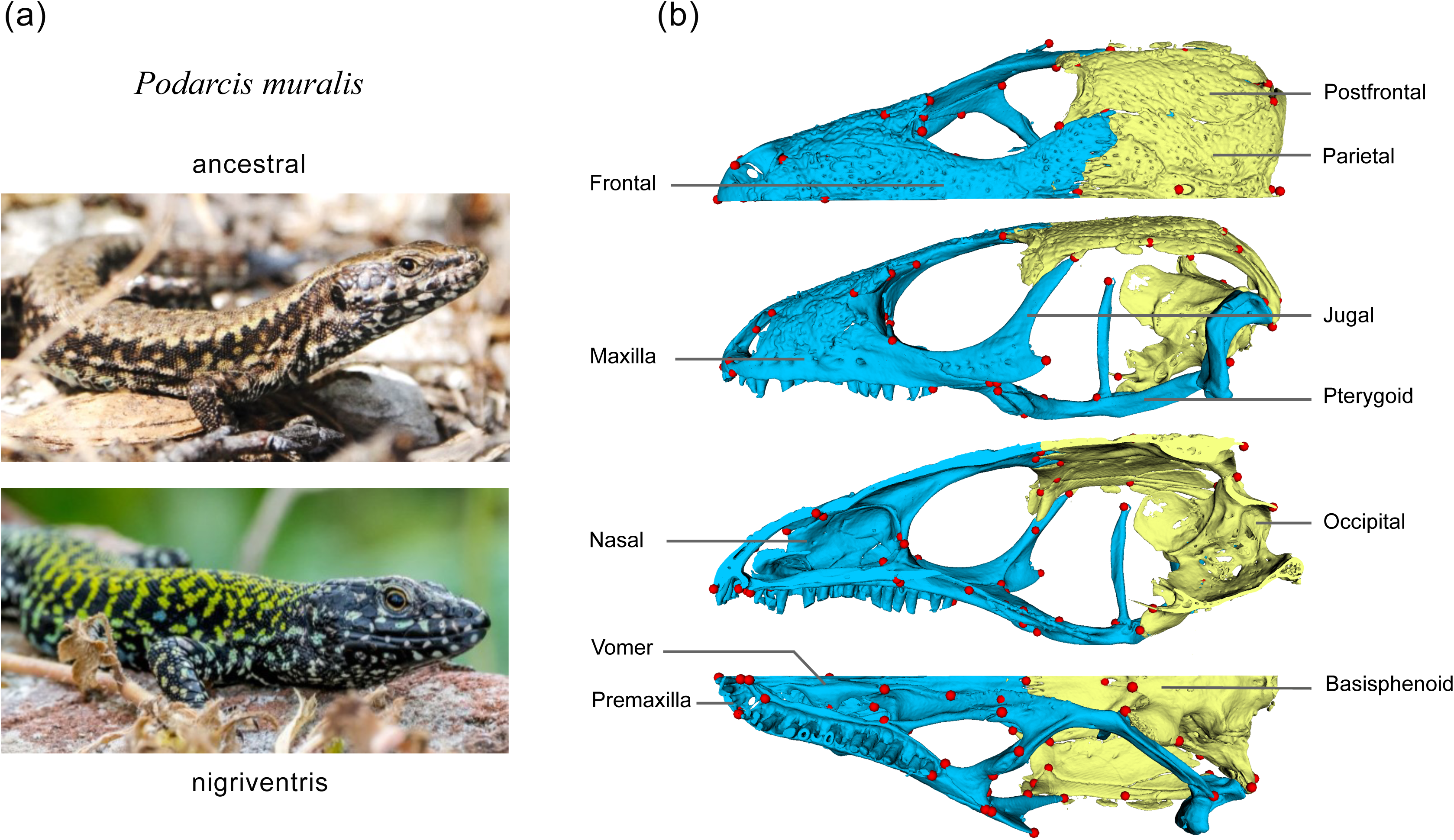
The nigriventris syndrome in *P. muralis* and skull anatomy in lacertids. (a) External appearance of males of the common wall lizards with the ancestral phenotype and the nigriventris syndrome. Top photograph by Nathalie Feiner, bottom photograph by Arnaud Badiane. (c) Skull partitioned into neural-crest (blue) and mesoderm-derived bones (yellow) in different perspectives (first: dorsal, second: lateral, third: sagittal, fourth: ventral). Red dots: landmark used in morphometric analyses.

Here, we used a comparative approach to test for a germ layer-associated developmental bias in the adaptive evolution of skull shape in lizards, using two levels of comparison. First, in the common wall lizard (*Podarcis muralis*), we investigated if changes in skull shape accompany the recent evolution of a sexually selected suite of traits, the ‘matricentric syndrome’, which consists of exaggerated coloration, morphological characters including larger relative head size, and aggressive behaviour (Figure 1; 19–21). The traits that comprise this syndrome partially originate from the neural crest, and genomic analyses suggest that genes that regulate aspects of neural crest cell biology contributed to its evolution. Thus, in *P. muralis*, we expect (i) modularity between mesoderm- and neural crest-derived bones, and (ii) that morphological change associated with the nigriventris syndrome is more pronounced for the parts of the skull with a neural crest origin.

Second, we assessed to what extent germ layer-associated developmental bias is maintained and contributes to morphological diversification of skulls within the lacertid lizards (family Lacertidae, Squamata). This family includes 387 currently described species (22; accession date 09/07/2025) distributed across Eurasia and Africa and their skull shape diversity has been linked to adaptation to diverse habitats and lifestyles (Hipsley & Müller, 2017). If germ layer-associated developmental bias shaped the diversification of skull morphology in this group, we expect (iii) that shape changes associated with the nigriventris phenotype in *P. muralis* provide explanatory power to the diversification of skull shape in lacertids, (iv) that modularity of mesoderm- and neural crest-derived bones persist across the lacertid family, and (v) that the tempo of evolution and disparity differ for mesoderm-*versus* neural crest-derived parts.

## Results

### Evolution of the nigriventris syndrome is associated with changes in skull size and shape

The nigriventris syndrome of the common wall lizard is composed of a suite of traits including coloration, morphology and behaviour that has been suggested to be associated with neural crest biology (Feiner *et al*., 2024). To test if skull shape differs between nigriventris and ancestral phenotypes, and if these differences are particularly pronounced for the parts of the skull that originate from the neural crest, we collected 3D scans of skulls of 62 specimens collected on the Italian peninsula using micro-computed tomography (µCT). We compared the skull shape of males from common wall lizards from populations classified along a 1 to 5 ordinal score, ranging from populations fixed for the ancestral phenotype (the ‘ancestral’ group, score 1), populations with intermediate phenotypes with scores 2 to 4 and populations fixed for the most extreme nigriventris phenotype (the ‘nigriventris’ group, score 5; Figure 1). For detailed analyses of the expression of the nigriventris syndrome across the Italian peninsula, see (Ruiz Miñano *et al*., 2021).

Consistent with previous findings (Feiner *et al*., 2024), common wall lizards from populations fixed for the nigriventris phenotype had longer heads relative to body size than specimens with the ancestral phenotype (Table S1). The ancestral and nigriventris phenotypes also differed in skull shape (Table 1), with individuals from populations with intermediate phenotypes tending to fall in between the two extremes (Table S1, Figure 2a). The inferred changes in skull shape from the ancestral to the derived nigriventris phenotype involved several neural crest-derived bones (jugal, epipterygoid and maxilla) being laterally compressed, thus giving a slimmer appearance to the skull. Other bones of neural crest origin tended to converge along the antero-posterior axis on the ventral side of the snout (epipterygoid, pterygoid, premaxilla, vomer) while being extended anteriorly along the dorsal side (frontal, nasal), resulting in relatively shorter snouts with smaller nares (Figure 2b). The changes in skull shape that accompanied the evolution of the nigriventris phenotype also involved mesoderm-derived bones, contributing to a lateral compression and a dorsal extension of the skull roof, giving the back of the skull roof a round aspect (parietal, post-frontal), and a relative contraction of the brain case (occipital bones and basisphenoid). Analyses based on shape estimates retaining size information led to the same conclusions (Figure S2). These results indicate that the nigriventris syndrome comprises head shape changes corresponding to a larger skull with elongated appearances and larger skull roofs relative to the snout.

**Figure 2.**
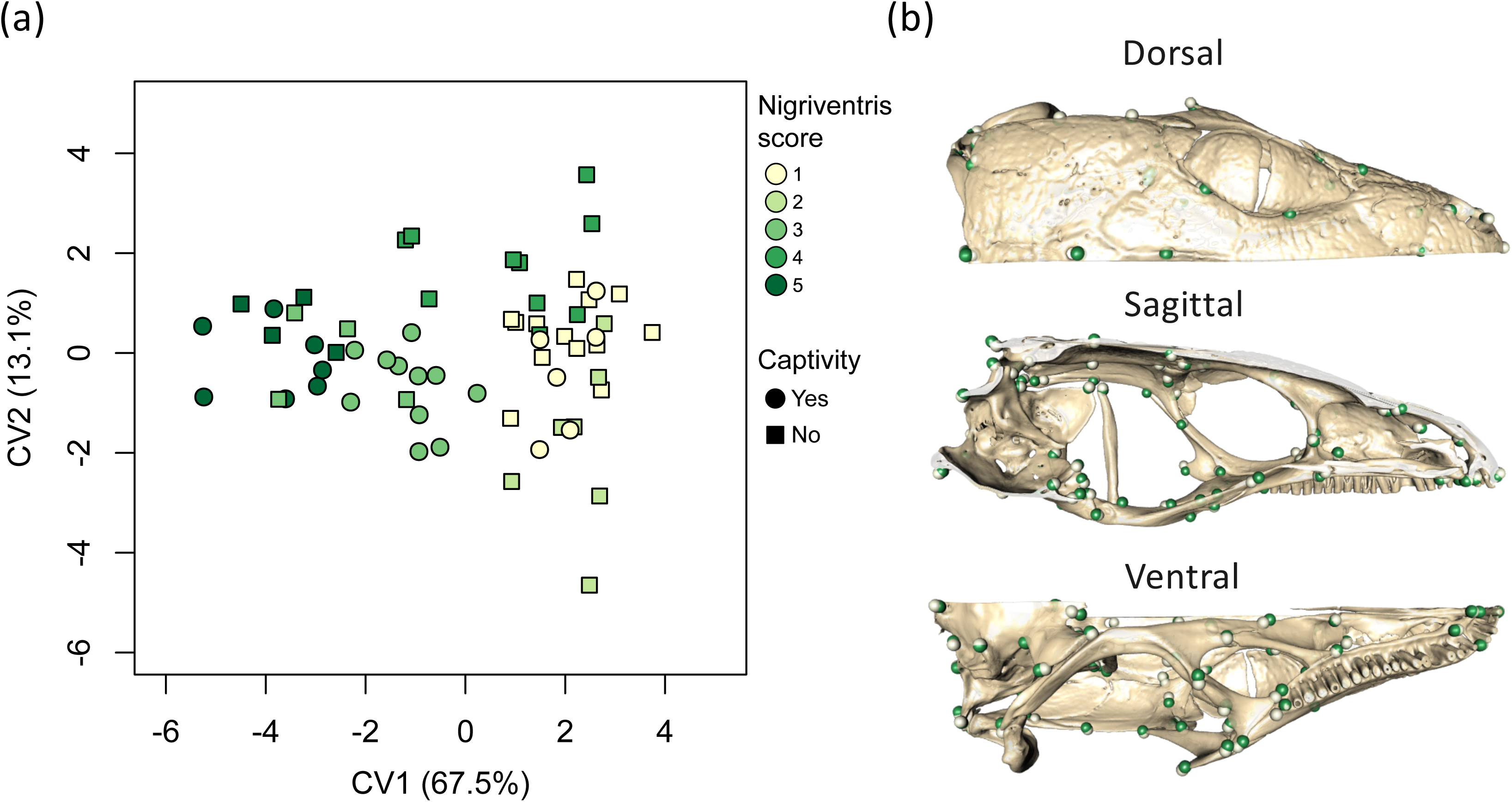
Canonical Variate analysis of landmark configurations on the skull of *Podarcis muralis*. Nigriventris score categories range from populations with ancestral-like phenotypes to populations with the most extreme expression of the nigriventris syndrome. (a) Projection of the first two canonical variates (CV), (b) landmark displacement between the two extremes of CV1 (minimum: dark green; maximum: light green).

**Table 1.** RRPP mANCOVA for shape changes in *P. muralis* varying in the expression of the nigriventris syndrome.

|  | **Df.** | **SS** | **MS** | ***R*<sup>2</sup>** | ***F*** | ***Z*** | ***P*** |
| --- | --- | --- | --- | --- | --- | --- | --- |
| Origin (captive, wild)<sup>1</sup> | 1 | 0.0030 | 0.0030 | 0.0455 | 3.326 | 3.741 | <0.001 |
| log(centroid size) | 1 | 0.0066 | 0.0066 | 0.1000 | 7.300 | 5.549 | <0.001 |
| nigriventris score | 4 | 0.0051 | 0.0013 | 0.0782 | 1.428 | 2.514 | 0.007 |
| log(centroid size) ×<br>nigriventris score | 4 | 0.0051 | 0.0013 | 0.0779 | 1.422 | 2.546 | 0.006 |
| Residuals | 51 | 0.0459 | 0.0009 | 0.6984 |  |  |  |
| Total | 61 | 0.066 |  |  |  |  |  |

<sup>1</sup>Fixed effect to correct for specimens that underwent to period of captivity following their capture in the wild.

### The dual origin of skull bones affects modularity and divergence within species

To test for co-variation between the mesoderm- and neural crest-derived elements of the skull, we partitioned the 77 landmarks by developmental origin (neural crest: 40; mesoderm: 35; uncertain: 2, see Methods) and assessed support for modularity within each configuration. We found that mesoderm- and neural crest-derived bones formed statistically supported modules in *P. muralis* (Covariance Ratio CR = 0.88; *Z* = −4.74; *P* < 0.001). Removing four landmarks that were on the parietal (a mesoderm-derived bone) but next to the intersection between the mesoderm- and neural crest-derived portions of the skull did not affect this result (CR = 0.89; *Z* = −3.89; *P* = 0.001). Pairwise comparisons of phenotypic distances between the ancestral and nigriventris group revealed significant and large effect sizes for the neural-crest derived module (*d* = 0.015; *Z* = 2.89; *P* = 0.002), while the mesoderm-derived module showed no statistically significant divergence between ancestral and nigriventris phenotypes (*d* = 0.012; *Z* = 0.83; *P* = 0.210). These results suggest that the germ layer origin structures skull shape variability in *P. muralis*, and that the neural crest-derived part of the skull is mostly responsible for the morphological changes associated with the nigriventris syndrome.

### Shape changes in the nigriventris phenotype provide explanatory power to the diversification of lacertid lizards

To assess the relationship between the divergence of ancestral and nigriventris skull shapes in *P. muralis* and macroevolutionary patterns of diversification, we first inspected how skull shape in lacertids varies along the dimensions that describe the differences between ancestral and nigriventris phenotypes. We collected data on skull shape for 174 species of lizards from the lacertid family, including species from all but one of the 43 genera (see Methods). When rotating the principal components of lacertid skulls along the axis of the maximum divergence between ancestral and nigriventris phenotypes, we find that 8.68% of the total variation in lacertids was captured by the ancestral-nigriventris axis of divergence (Figure 3). In comparison, the first three principal components capture 26.17%, 17.17% and 6.20% of the total variation. Species with extreme values in the direction of the nigriventris phenotype are lizards with robust skulls, relatively small nares and orbits, and large parietal bones, like large-bodied species from the *Timon*, *Gastropholis*, *Gallotia* and *Lacerta* genera, but also the smaller forest dwelling species *Adolphus jacksoni*. Species with extreme values towards the ancestral shape of *P. muralis* possess slender snouts, large dorsally exposed nares and large orbits, reminiscent of a previously described paedomorphic appearance in some lacertids (Hipsley & Müller, 2017). These species include small-bodied, sand or desert dwelling species (genera *Acanthodactylus*, *Heliobolus*, *Meroles* and *Mesalina*) as well as the tree gliding *Holaspis laevis*. We conclude that the divergence between ancestral and nigriventris morphotypes in *P. muralis* spans an axis that describes a considerable amount of variation in skull shape across lacertid lizards.

**Figure 3.**
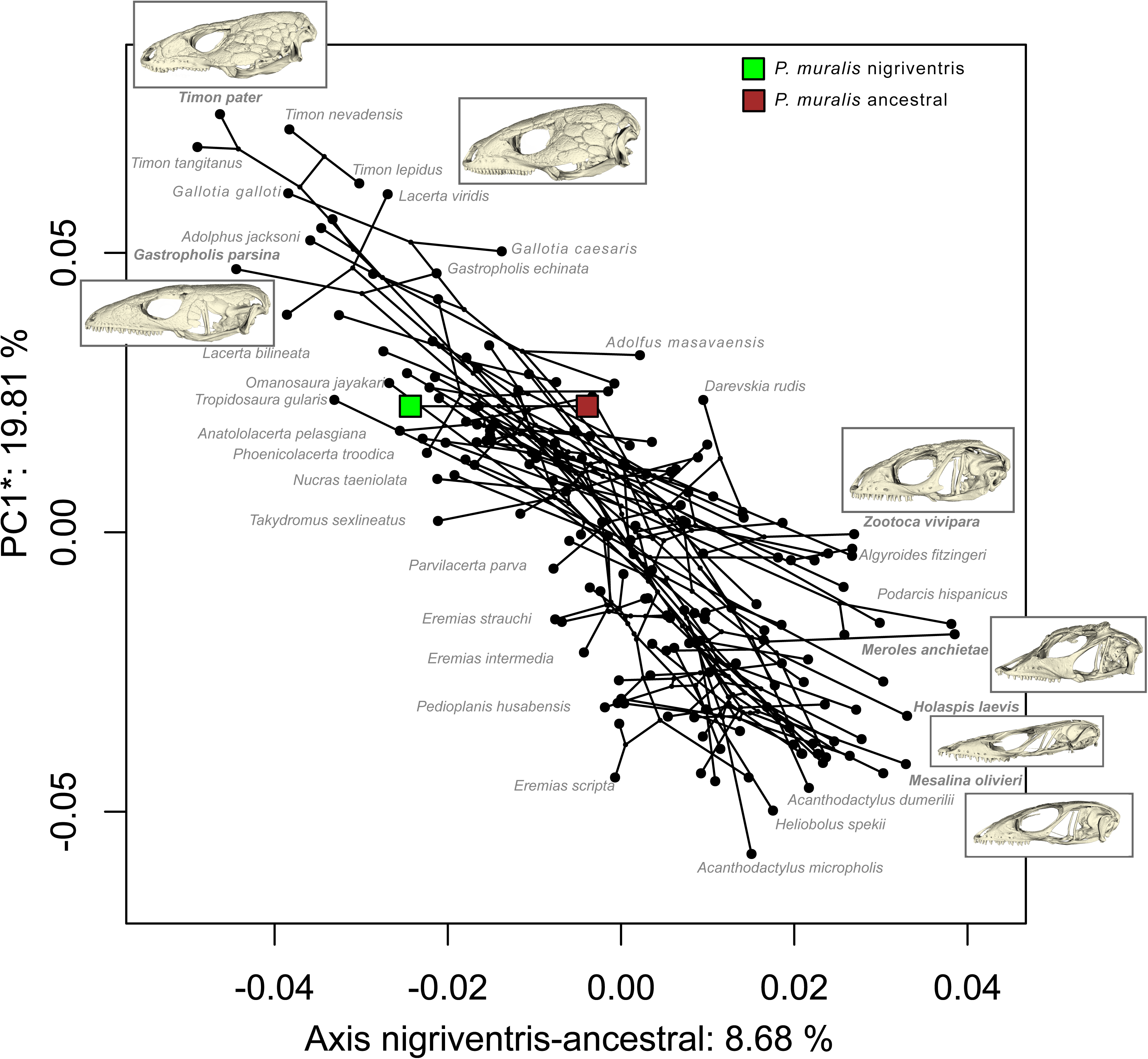
Morphological variation in lacertids with regards to the ancestral-nigriventris axis of divergence. Phylomorphospace of 174 species of lacertids (black dots) aligned with the axis of divergence between the ancestral and the nigriventris average shapes in *P. muralis* (shown as brown and green squares). The y-axis is the first principal component minus the variation from the rotated axis. To provide examples of the morphologies at the extreme ends of the ancestral-nigriventris axis, we show skull morphologies of selected species (highlighted in bold text).

### Germ layer-associated modularity persists at the macroevolutionary scale

If the contrasting variational properties induced by the dual developmental origin have a persistent effect on diversification beyond the species level, we expect modularity between the mesoderm- and neural crest-derived regions of the skull in lacertids, in agreement with our observations in *P. muralis*. Further, we expect differences between modules in their evolutionary rates and in morphological variance (i.e., disparity). Despite 50-70 million years of adaptive divergence to meet a diversity of functional demands (Hipsley *et al*., 2009; Čerňanský *et al*., 2020), patterns of covariation between skull bones are consistent: as for *P. muralis*, configurations of mesoderm- and neural crest-derived bones formed statistically supported modules within lacertids (CR = 0.967; *Z* = −4.15; *P* = 0.002). These results suggest that the dual origin of the lacertid skull structures morphological variation at the population level within *P. muralis* as well as in deep evolutionary time, at the level of the lacertid family.

Considering the tempo of evolution for the two modules, we find that the neural crest-derived module shows an overall slower evolutionary rate relative to the mesoderm-derived module (σ^2^_mult neural crest_ = 5.11×10^−6^; σ^2^ = 7.95×10^−6^; *Z* = 11.98; *P* < 0.001). Consistent with this, phylogenetic signal was higher for the neural crest-relative to the mesoderm-derived module (*Z*_neural crest_ = 28.63; *Z*_mesoderm_ = 23.73; pairwise *Z* = 28.63; *P* < 0.001). We also find that the neural crest-derived module overall showed reduced disparity (Procrustes variance, PV, neural crest module: 0.0017; PV mesoderm module: 0.0023; average per-landmark variance: neural crest = 4.32×10^−5^; mesoderm = 6.47×10^−5^; *t* = 3.23; d.f. = 64.68; P = 0.002). Thus, in contrast to the morphological divergence involved in the nigriventris syndrome in *P. muralis*, the neural crest-derived part of the skull shows lower realized evolvability relative to the mesoderm-derived part at the level of the family, both in terms of evolutionary rate and disparity.

Finally, we tested at the landmark-level whether the specific regions of the skull that showed high divergence between ancestral and nigriventris phenotypes in *P. muralis* exhibit particularly high variation and evolutionary rates across species. We found that landmarks describing the largest phenotypic distances between the ancestral and the nigriventris phenotypes in *P. muralis* were indeed associated with higher disparity (i.e., per-landmark variance) within lacertids (Figure 4a, adjusted R² = 0.23). However, this effect was stronger for landmarks placed on mesoderm-than neural crest-derived bones, as indicated a tendency for differences between the slopes estimates of each type of landmark (slope estimates [95% Confidence Intervals]: β_neural crest_ = 0.004 [−0.007; 0.016]; β_mesoderm_ = 0.015 [0.006; 0.024]; *t* = −1.82; *d.f.* = 71; *P* = 0.072). Furthermore, while we observed an overall positive relationship between per-landmark evolutionary rates and their associated phenotypic distances between the ancestral and nigriventris phenotypes (Figure 4b; adjusted R² = 0.18), comparisons of slopes suggested that this effect was driven by mesoderm-derived bones (β_neural crest_ > 0.001 [−0.005; 0.006]; β_mesoderm_ = 0.006 [0.002; 0.010]; *t*-test: *d.f.* = 71; *t* = −2.11; *P* = 0.038). One interpretation of this pattern is that microevolutionary divergence in the neural crest portion of the skull may be largely uncoupled from patterns of disparity and evolutionary rates at the macroevolutionary level.

**Figure 4.**
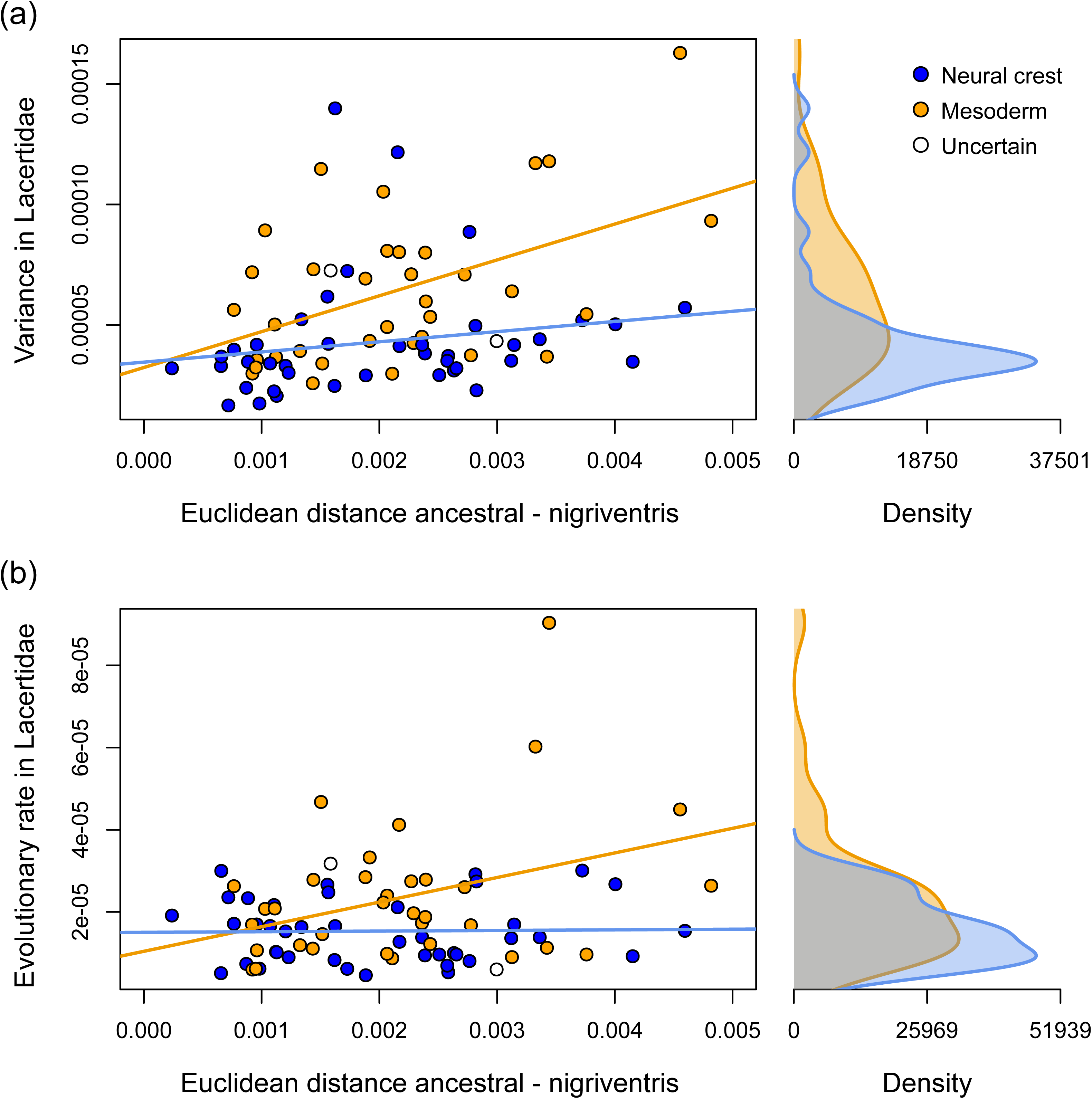
Relationship between per-landmark magnitude of change in the nigriventris phenotype and the variation at the macroevolutionary level. Kernel densities of variances and evolutionary rates of the two types of landmarks shown on the right insets. (a) Per-landmark Euclidean distances between the ancestral versus nigriventris groups over per-landmark variances in lacertids (adjusted *R²* = 0.22). (b) Per-landmark Euclidean distances between the ancestral vs nigriventris groups over per-landmark evolutionary rates in lacertids (adjusted *R²* = 0.18).

Altogether, these results suggest that while the neural crest-derived part of the skull contributed disproportionally to the evolution of the nigriventris syndrome within *P. muralis*, the neural crest-derived part is the least evolvable part of the skull in lacertids. That the parts of the skull derived from the neural crest is highly conserved but can be associated with rapid phenotypic divergence is supported by the observation that rare and localised shifts in the evolvability of the neural crest-derived module accumulate in the genus *Podarcis* (Figure 5). However, we did not find support for an overall faster rate of skull evolution in *Podarcis* relative to other lacertids (σ²*_Podarcis_* = 1.25×10^−6^; σ²_other lacertids_ = 7.24×10^−6^; *Z* = 0.13; *P* = 0.49).

**Figure 5.**
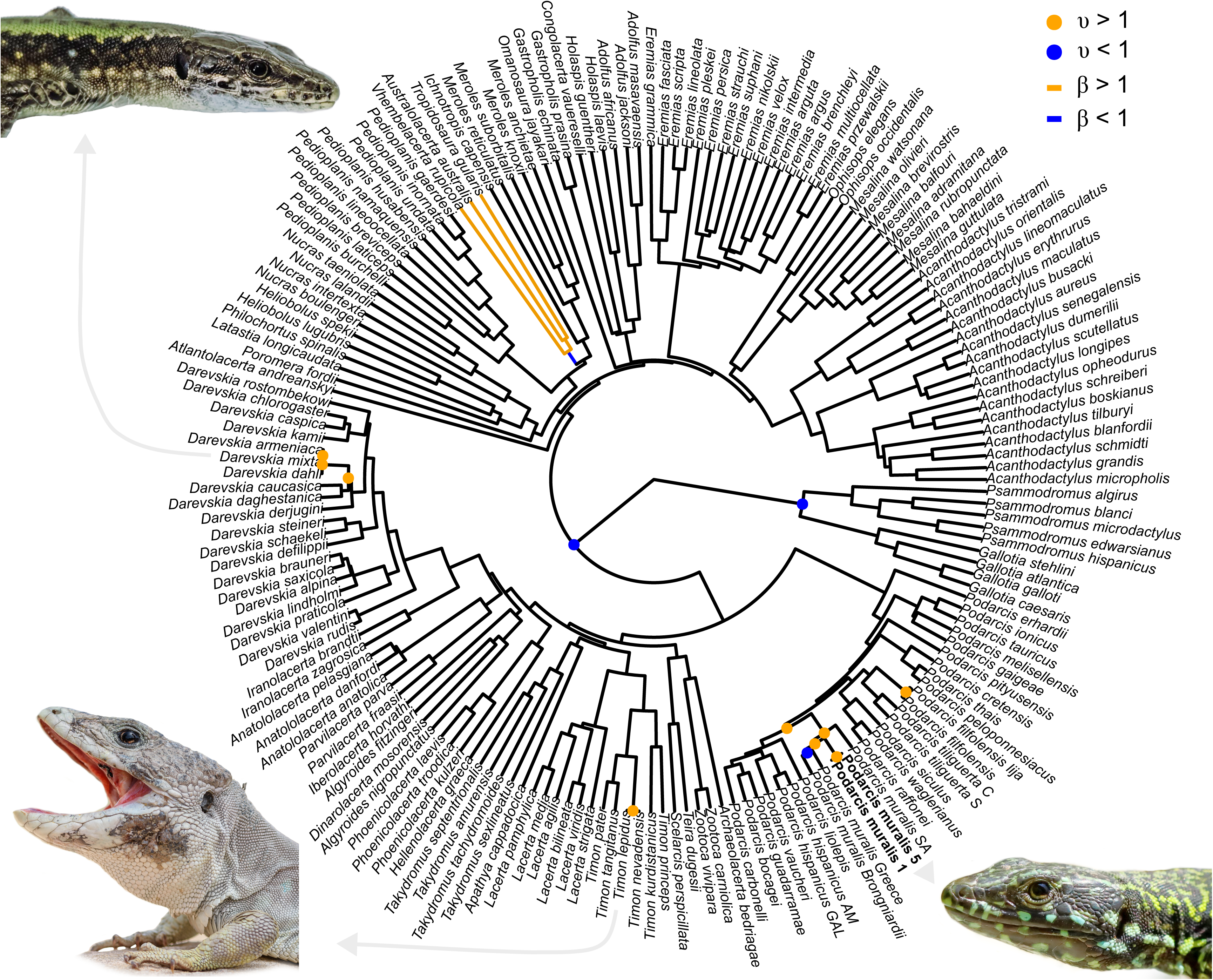
Shifts in the evolvability of the neural crest-derived part of the skull across the phylogeny of lacertids. Shifts in evolvability (υ) and in selection (β) were obtained with phylogenetic modelling (Fabric model), using the first five principal components of the phylomorphospace of the landmarks placed on the neural crest-derived bones. Only the shifts with posterior probability > 0.8 are shown. The Central Italy lineage of *P. muralis* features two tips, one for each average shape of the nigriventris scores 1 (ancestral phenotype) and 5 (most pronounced nigriventris phenotype). Insets show representative males of species within clades where shifts in evolvability occur. Photographs by Giorgi Iankoshvili (*D. mixta*), Pedro Luna Guillen (*T. nevadensis*) and Quentin Horta-Lacueva (*P. muralis*, nigriventris phenotype).

## Discussion

In this study, we show that the evolution of the vertebrate skull is shaped by the developmental origins of its bones, which arise from either migratory neural crest cells or mesodermal tissue. Skull shape variation within and among species of lacertid lizards is partitioned into two distinct modules corresponding to these developmental origins. The contributions of each module to phenotypic variation differ across evolutionary time scales. At the microevolutionary level, among common wall lizards, the neural crest-derived module accounts for most of the cranial variation associated with the sexually selected nigriventris syndrome. In contrast, at the level of the family Lacertidae, this same module exhibits lower evolutionary rates and reduced morphological disparity. We propose that these contrasting patterns can be explained by the neural crest’s dual role: while it can enable rapid response to correlational sexual selection on coloration, head morphology and behaviour, such pleiotropy also limits its role for the adaptive divergence of skull shape on a macroevolutionary scale.

The capacity of the neural crest to structure morphological variability at the species level has been most extensively explored within the framework of the domestication syndrome. Correlated changes in coloration, behaviour and craniofacial morphology observed in domesticated animals have been hypothesized to arise from their shared developmental origin in neural crest cells (Wilkins *et al*., 2014; Sánchez-Villagra *et al*., 2016). Support for the neural crest’s involvement in domestication remains equivocal (Pendleton *et al*., 2018; Johnsson *et al*., 2021; Wilkins *et al*., 2021; Rubio & Summers, 2022; Le Verger *et al*., 2024). For example, a lack of modularity among mesoderm- and neural crest-derived bones in chickens suggests that neural crest cells can play a limited role for the variability in the skull shape of domesticated animals (Stange *et al*., 2018). Given that signatures of domestication events may have been superseded by subsequent selection for extreme phenotypes (Sánchez-Villagra *et al*., 2017), the hypothesis may apply more directly to wild populations (Brandon *et al*., 2023). Our previous work has identified the evolution of the sexually selected nigriventris syndrome in *Podarcis muralis* as a promising case study. Genomic regions associated with this syndrome are enriched for genes with a described function in neural crest cell regulation (Feiner *et al*., 2024), and coloration, morphology and behaviour remain integrated during introgression into a distantly related lineage (While *et al*., 2015; Feiner *et al*., 2024). Here, we report that the neural crest-derived module contributed more to the divergence between ancestral and nigriventris phenotypes than the mesoderm derived module. This finding supports the hypothesis that the neural crest plays a pivotal role in the evolution of the nigriventris syndrome and provides evidence that morphological variation generated by neural crest cells can shape evolutionary trajectories in natural populations.

At the macroevolutionary scale, studies on placental mammals and birds have shown that bones derived from the neural crest show higher evolutionary rates than mesoderm-derived bones (Felice & Goswami, 2018; Goswami *et al*., 2023). While we also found that the neural crest-derived bones form a module, it displayed slower evolutionary rates and reduced morphological disparity, as well as stronger phylogenetic signal. Furthermore, the neural crest-derived landmarks that contributed most significantly to the nigriventris syndrome did not exhibit accelerated rates of evolution, indicating that the neural crest-derived module does not contribute disproportionally to skull shape diversification at a broad phylogenetic scale. How such evolutionary rates in the neural crest may affect evolvability and evolutionary trajectories is difficult to establish. In placentals, mesoderm-derived bones are as prone as neural crest-derived bones to generate disparity despite their slower evolution (Goswami *et al*., 2023). That our results contrast with the patterns observed in other clades, and differ between micro- and macroevolutionary scales within lacertid lizards, indicates that even though development structures variability, the realised variation within each portion of the skull can be complex to predict.

A possible explanation for the greater importance of the mesoderm in the diversification of the lacertid skull is that functional morphology related to feeding performance and ecological specialization could depends more heavily on mesoderm-derived skeletal elements. However, this idea is not supported by studies on *Podarcis melisellensis* showing that the shapes of several neural crest-derived bones (e.g., pterygoid, jugal, premaxilla) covary with diet and bite force (Taverne *et al*., 2021). Stronger and bigger head muscles were described as being associated with shorter snouts as well as taller skulls, likely from the increased curvature of the pterygoid, a neural crest-derived bone (Taverne *et al*., 2021). Furthermore, skull kinesis in lizards requires articulated movements between bones of both mesoderm and neural crest origins (Gans *et al*., 2008), suggesting that biomechanical factors may play a limited role in explaining germ layer-associated differences in skull diversification. Instead, the mesoderm-derived part of the skull may show greater evolvability since several of these bones ossify relatively late in ontogeny (Barahona & Barbadillo, 1998). Late ossification has been proposed to explain the evolution of more prominent posterior parts of the skull in males of *Podarcis bocagei* and *P. carbonelli* (Kaliontzopoulou *et al*., 2008), and might confer higher variability to the mesoderm-derived part of the skull across lacertids.

The lower disparity observed in neural crest-derived regions across lacertid lizards may give the impression that the rapid cranial divergence associated with the nigriventris syndrome reflects a release from developmental constraint. However, we suggest that the development of head morphology, coloration, and behaviour in fact makes neural crest cells facilitate coordinated responses to correlational sexual selection. In *Podarcis muralis*, this is supported by evidence of selection acting on multiple sexually selected traits (e.g. coloration and relative head size), many of which partially originate from the neural crest (While *et al*., 2015; MacGregor *et al*., 2017b). In contrast, cranial adaptations reflecting ecological specialization and diet could be readily achieved by the mesoderm-derived module being potentially less pleiotropic and more developmentally flexible (Kaliontzopoulou *et al*., 2008; Brandon *et al*., 2023), thereby driving the ecological specialization that is responsible for much of the diversity of lacertid skull shape. Yet, the axis of divergence associated with the nigriventris phenotype does align with one of the major directions of morphological variation in lacertids. Moreover, the skull regions contributing most to the evolution of the nigriventris syndrome also exhibit elevated variability and evolutionary rates across species. This suggests that the nigriventris phenotype evolved along a major axis of cranial variation shared within lacertids, with sexual selection acting as a key driver of divergence in morphology along this axis. Consistent with this view, shifts in evolvability of neural crest-derived regions appear to have occurred independently in other lacertid lineages: in other *Podarcis* species, in *Timon* species with pronounced sexual dimorphism (*T. nevadensis*), and in *Darevskia*, which includes parthenogenetic, all-female species (41, see 42 on pathenogenesis and head shape in *Darevskia*). Together, these patterns suggest that changes in correlational sexual selection may repeatedly shape the evolvability of neural crest-derived traits across lacertids.

The contrasting modular patterns of skull variation across evolutionary levels provide insights on the relationship between population-level trait variabilities and long-term evolutionary trends. Several recent studies suggest that macroevolution is predictable from how traits covary (Houle *et al*., 2017; Holstad *et al*., 2024; Schluter, 2024; Tsuboi *et al*., 2024; Rohner & Berger, 2025). However, while the consistency of correlational structures imposed by the dual developmental origin of the skull supports this view, our results illustrate why a persistent developmental bias need not to result in alignment between major axes of variation within and among species.

## Methods

### Data collection

We studied skull morphology using high-resolution, μCT scans of the skulls of 263 ethanol-preserved specimens. Scanning was performed on SkyScan1173 and FF20 CT – Comet Yxlon scanners (voxel sizes 7.45-28.04μm), image stacks were reconstructed with NRecon and CERA, the resulting volumes were segmented with manual thresholding and editing tools before being converted to meshes, and decimated using 3D Slicer (Fedorov *et al*., 2012) and SlicerMorph (Rolfe *et al*., 2021). When overly rugged surfaces were observed by visual inspection, surfaces were smoothened.

The data contained two subsets of specimens dedicated to micro- and macroevolutionary questions. One subset contained 62 adult male *Podarcis muralis* (the “microevolutionary data”). The specimens belonged to the Central Italy lineage (Yang *et al*., 2022), in which the nigriventris phenotype originated. The *P. muralis* specimens belonged to collections from Museum Koenig Bonn (*n* = 30) and Natural History Museum Vienna (*n* = 8), and from captive maintenance of wild-caught individuals at Lund University (Pranter & Feiner, 2025) (*n* = 24). The expression of the nigriventris syndrome could not be quantified directly on museum specimens because the ethanol-preservation alters pigmentation. Therefore, we attributed an ordinal score to every specimen by mapping their capture locality to the phenotypic gradient of the nigriventris syndrome established in previous studies (16, 17, Figure S1). That way, we produced a nigriventris score ranging from 1 (no exaggerated trait, “ancestral” phenotype) to 5 (most extreme values of the syndrome of exaggerated traits, typical “nigriventris” phenotype).

The other subset contained one specimen for each of 174 species and major lineages with unresolved taxonomic status from the family Lacertidae (the “macroevolutionary data”, ∼44% of the described species, 42 of the 43 genera, the genus with the single species *Dalmatolcacerta oxycephala* missing). We considered as major lineages: the Corsican and Sardinian lineages of *Podarcis tiliguerta* (Salvi *et al*., 2017), three lineages of *Podarcis muralis* described as “Southern Alps”, “Southern Balkan” and “Western Europe” (Yang *et al*., 2022), and two lineages of *P. filfolensis* from Malta and Linosa (Salvi *et al*., 2014), and the “Galera” and “Albacete/Murcia” lineages of *P. hispanicus* (Kaliontzopoulou *et al*., 2011; Yang *et al*., 2021). All specimens were males except for three parthenogenetic, female-only species (*Darevskia rostombekowi*, *D. armeniaca* and *D. dahli*).

For the macroevolutionary dataset, we generated μCT scans of 147 specimens, to which we added 25 previously scanned specimens from Hipsley and Müller (Hipsley & Müller, 2017) and two scanned specimens from MorphoSource (UF:Herp:130038 ark:/87602/m4/486159; UF:Herp:91747 ark:/87602/m4/604781).

The phylogenetic analyses relied on the ultrametric tree from Title and colleagues (Title *et al*., 2024). We used *phytools* (Revell, 2012) to preprocess the phylogenetic data and append the tree with additional species or major lineages according to the divergence times given in the following references: *Acanthodactylus erythrurus lineomaculatus* (Rancilhac *et al*., 2023), *Eremias fasciata* (Orlova *et al*., 2023), *Gastropholis echinata* (Tonini *et al*., 2016), *P. thais* (Kiourtsoglou *et al*., 2021), *P. ionicus* (Psonis *et al*., 2021), *Zootoca carniolica* (Cornetti *et al*., 2015) and the major lineages of *P. filfolensis, P. hispanicus, P. muralis* and *P. tiliguerta* (Salvi *et al*., 2014).

### Geometric morphometrics

We digitized 77 landmarks on the right side of skull models of every specimen with the Markups module in 3D Slicer 5.2.1. (Fedorov *et al*., 2012). In cases where bones were deformed or broken, we mirrored the model and digitized what was originally the left side. The landmark data were imported into R with SlicerMorphR (Rolfe *et al*., 2021). All landmark configurations were then superimposed to a common coordinate system through Generalised Procrustes Analyses (GPA) in *geomorph* (Adams & Otárola-Castillo, 2013; Adams *et al*., 2013). We conducted the GPA after producing two-sided landmark configurations by mirroring the right-side to avoid inflated landmark variations along the midline (Bardua *et al*., 2019). The artificially produced left-side was discarded after the GPA, and the analyses were conducted on the aligned configuration of the original side.

### Microevolution 1: Shape changes in P. muralis as part of the nigriventris syndrome

We first tested for differences in skull shape within *P. muralis* based on ordinal scores for the nigriventris syndrome. We performed np-MANCOVA using Randomized Residual Permutation Procedure (RRPP) with 10 000 permutations and type I sum of squares (Adams & Otárola-Castillo, 2013). We accounted for the effects of the extended periods of captivity, centroid size, and from potential variation in allometry among ordinal categories by adding fixed effect in sequential order as in the full model:

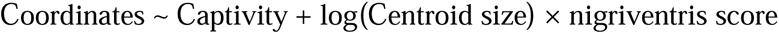

Although our main model suggested unique allometries among groups (Table1), pairwise comparisons returned statistical support for allometric differences only between the ancestral group and the three groups of intermediate categories (Table S2). We therefore considered it appropriate to use size-regressed residuals of shape changes for downstream analyses of differences between the ancestral versus nigriventris phenotype.

Finally, we studied the patterns of shape changes among ordinal categories of the nigriventris syndrome by conducting Canonical Variate Analyses (CVA) adapted to multidimensional data, using *Morpho* (Schlager, 2017). We visualized the 3D changes in landmark configuration along the covariate axes as a deformation of the cubic grid (thin-plate spline interpolation), using *Morpho* and *rgl* (Adler *et al*., 2020).

### Microevolution 2: Importance of the neural crest in the nigriventris-associated shape variation

We tested whether variational properties of neural crest-derived bones (NC) relative to mesoderm-derived bones (MES) explained the skull shape changes observed in *P. muralis.* We proceeded in two steps as follows. First, we conducted modularity analyses to test for separate patterns of shape variation related to either tissue type. We partitioned the landmark data into two subsets according to their germ layer of origin: NC bones (40 landmarks), MES bones (35 landmarks). We attributed a NC or MES origin to each bone by relying on fate-maps of model vertebrates (Maddin *et al*., 2016) and on studies of homology in extant and extinct taxa (Fabbri *et al*., 2017; Smith-Paredes *et al*., 2018). Two landmarks placed on the squamosal, for which we found no information of its developmental origins, were labelled as uncertain. We evaluated modularity between the two sections with Covariance Ratio (CR) coefficients obtained via permutation (Adams & Collyer, 2016).

Since we observe modularity between neural crest- and mesoderm-derived bones (see Results), we tested whether the NC module explained shape changes associated with the nigriventris syndrome. We proceeded by reanalysing head shape variations based on ordinal scores with RRPP MANCOVA (see *Microevolution 1*), this time only considering the landmark either from the NC or from the MES module. We performed pairwise group comparisons between the two most extreme ordinal categories (“ancestral” or “nigriventris”) and assess the effect size and significance of the differences between groups for each module.

### Macroevolution 1: Importance of the neural crest in the evolution of lacertids

We tested for modularity in lacertids by estimating Covariance Ratio coefficients (*CR*) between the NC and MES modules that were defined on the same subset of landmarks as for *P. muralis*. Furthermore, we tested for differences in phylogenetic signal, evolutionary rates and disparity between the two modules by comparing *Z* estimates (Collyer *et al*., 2022) and multivariate evolutionary rate ratios *R* (Denton & Adams, 2015), respectively.

We visualized morphological diversity in lacertids in relation to the nigriventris phenotype by rotating the phylomorphospace (rigid rotation) along the vector describing shape changes between the two extreme ordinal nigriventris categories in *P. muralis*. To test whether the anatomical regions responsible for shape changes for the nigriventris syndrome in *P. muralis* were responsible for higher variation and evolutionary rates in lacertids, we estimated the magnitude of landmark changes associated with the nigriventris syndrome by calculating the Euclidean distances of each homologous pair of landmarks between the mean landmark configuration of the “ancestral” (Ordinal score 1) and the “nigriventris” group (Ordinal score 5) in the *P. muralis* data. We evaluated per-landmark variances and per-landmark evolutionary rates in lacertids by following Fabre and colleagues (Fabre *et al*., 2020). Finally, we assessed the effect of the NC and the MES origin in the relationship between per-landmark effect sizes in the nigriventris phenotype and per-landmark variances or evolutionary rates in lacertids by using linear regressions. Note that highly influential landmarks had little effect on these results (e.g., Cook’s distances of all landmarks were below 1, and the removal of a landmark with relatively high Cook’s distance did not change the results of either regression).

We investigated the possibility of shifts in evolvability in the NC by performing phylogenetic modelling with the Fabric model in BayesTrait (Pagel *et al*., 2022). For the NC and MES modules, we used the first five and four principal components of the phylomorphospace as shape variables (the subsequent components individually explained 4% of the variance or less), which described 57.8% and 65.9% of the total variation, respectively. Model convergence was estimated based on trace and posterior density plots.

## Data and Code availability

Landmark data, skull models and metadata are available on Zenodo: https://zenodo.org/records/15855204?preview=1&token=eyJhbGciOiJIUzUxMiJ9.eyJpZCI6IjFl <u>MTQyYWE1LTRiYWMtNGRhMC1iMzcyLTA0NWIwZWZkNjg5NCIsImRhdGEiOnt9LCJyY W5kb20iOiI5YWNhM2UxOTQ5YWQzM2UwODNkZjRkYTNmN2IyNDQzMCJ9.04Rcf3p5A 9bznlKOH8BfQG3RI4LJ3jA_wxcn1WpbaEHcrlkqf5vuf0Kq5iIxhIwSwQVFmmrzx4BEbJu_dB 3Rdg</u>

Scripts are available on Github: https://github.com/quentin-evo/lacertid-skull-NC

## Author Contributions

QH, NF, TU designed the research; QH performed the research; QH preformed the analyses; QH, NF, TU interpreted the data; QH, NF, TU wrote the paper; CH, MG, MF, JM, MK contributed data; CH, MG, MF, JM critically revised the manuscript.

## Acknowledgment.

We are grateful to Benjamin Wipfler and Juliane Vehof (Museum Koenig Bonn), to Karolin Engelkes and Lena Schwinger (Museum der Nature Hamburg), to Franck Tillack (Museum für Naturkunde Berlin) and to Ola Gustafsson (Microscopy Facility at the Department of Biology, Lund University) for providing access to museum collections and CT-scanning facilities, and for providing support during scanning and data collection. Specimens from Naturhistorisches Museum Wien were loaned thanks to the contribution of Georg Gassner. We thank Coleman Sheehy and David Blackburn (Florida Museum of Natural History) for providing on specimens downloaded from Morphosource. We thank Javier Abalos, Lisandro Milocco and members of the Feiner-Uller group (Lund University) for their input during data analyses and the interpretations of the results.

