## Supplementary material for "Neural crest cell biology shapes lizard skull evolution across evolutionary time scales": Figure S1, Figure S2, Table S1

### Supplementary Figures and Tables

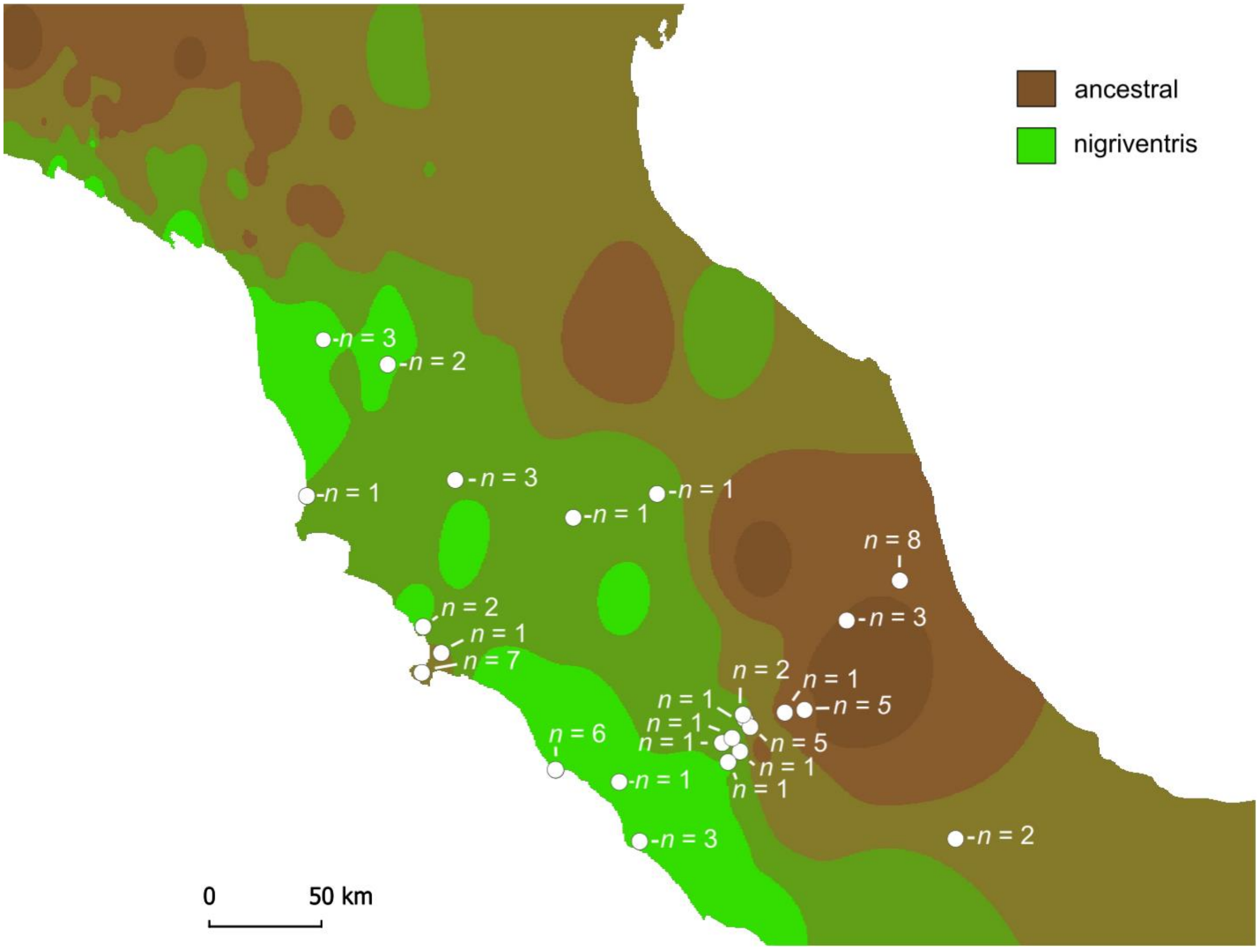


**Fig. S1.** Locations and sample sizes of *Podarcis muralis* specimens from central Italy used for the study. Distribution of dorsal green coloration, a proxy of the strength of the nigriventris syndrome, from ancestral type to extreme nigriventris phenotype, is based on the estimates from (1, 2). The dorsal green colour variations distribution is coarsely extrapolated for illustrative purpose, the final decisions to attribute nigriventris scores to the populations were made based on thorough considerations of the fine scale data from (1, 2).

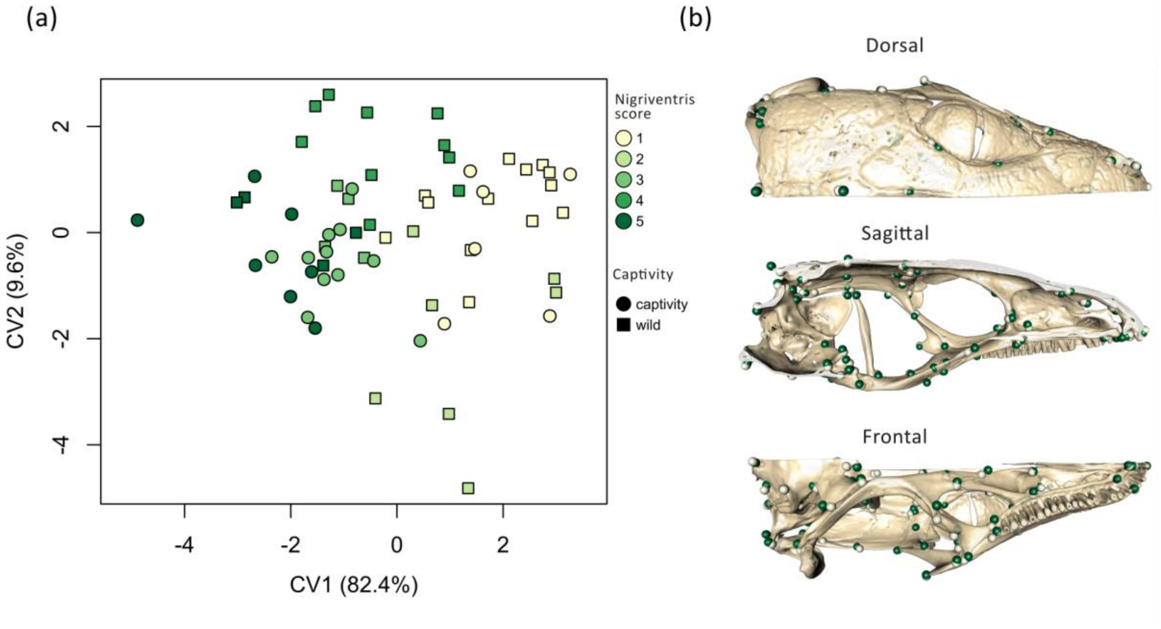


**Fig. S2.** Canonical variate analysis of landmark configurations on the skull of *Podarcis muralis* with raw data instead of size-corrected residuals. Nigriventris score categories range from populations with ancestral like phenotype to populations with the most extreme expressions of the nigriventris syndrome. (a) Projection of the first two canonical variates (CV), (b) landmark displacement between the two extremes of CV1 (min: light green; max: dark green). Inverse of CV1 displayed for comparisons with Figure 2.

**Table S1.** Posterior estimates of head length (linear measurements) in *P. muralis* from a linear model (MCMC GLMM) including nigriventris score as a fixed effect [1].

| Effect | Posterior mode | 95% CrI | Effective sample size | pMCMC |
|---|---|---|---|---|
| Origin - captivity | 0.271 | -0.066: 0.63 | 1000 | 0.13 |
| Origin - field | -0.26 | -0.439: -0.098 | 1000 | 0.004 |
| nigriventris score 2 | 0.169 | -0.835: 1.277 | 1000 | 0.636 |
| nigriventris score 3 | 0.713 | -0.644: 1.498 | 1000 | 0.482 |
| nigriventris score 4 | 0.336 | -0.233: 0.906 | 1251 | 0.334 |
| nigriventris score 5 | 0.337 | **0.023: 0.467** | 1000 | **0.016** |
| Body length (SVL) | 0.245 | 0.198: 0.268 | 1000 | 0.001 |
| nigriventris score 2 x SVL | -0.059 | -0.187: 0.039 | 1000 | 0.228 |
| nigriventris score 3 x SVL | 0.103 | -0.087: 0.226 | 1000 | 0.296 |
| nigriventris score 4 x SVL | 0.027 | -0.057: 0.095 | 1000 | 0.594 |
| nigriventris score 5 x SVL | 0.021 | -0.017: 0.068 | 1000 | 0.302 |

[1]Formula: head length ~ captivity + nigriventris score x body length – 1. Iterations: 130,000; thinning burnins: 30,000; thinning intervals: 100. Baseline: nigriventris score 1.

**Table S2.** Pairwise comparison of the allometric slope of nigriventris categories in *P. muralis*.

| | Procrustes distances | | | Vector correlations | | | |
|---|---|---|---|---|---|---|---|
| Pairs | *d* | Z | $P_{distance}$ | *r* | angle | Z | $P_{angle}$ |
| 1 vs. 2 | 0.022 | **1.810** | **0.036** | -0.15 | 1.721 | **3.546** | **<0.001** |
| 1 vs. 3 | 0.016 | **2.171** | **0.014** | 0.281 | 1.286 | **2.639** | **0.004** |
| 1 vs. 4 | 0.013 | 0.320 | 0.374 | 0.264 | 1.303 | **3.328** | **<0.001** |
| 1 vs. 5 | 0.019 | **2.042** | **0.019** | 0.46 | 1.092 | 0.902 | 0.184 |
| 2 vs. 3 | 0.021 | 0.865 | 0.192 | 0.375 | 1.186 | 0.276 | 0.39 |
| 2 vs. 4 | 0.021 | 1.476 | 0.068 | 0.519 | 1.025 | -0.794 | 0.789 |
| 2 vs. 5 | 0.022 | 0.488 | 0.315 | 0.169 | 1.401 | 1.498 | 0.066 |
| 3 vs. 4 | 0.014 | -0.992 | 0.840 | 0.728 | 0.755 | -0.735 | 0.768 |
| 3 vs. 5 | 0.016 | 0.862 | 0.195 | 0.485 | 1.064 | 0.768 | 0.221 |
| 4 vs. 5 | 0.016 | -0.788 | 0.786 | 0.56 | 0.977 | 0.684 | 0.244 |
